# Desynchronized Somas and Terminals in a Morning Clock Neuron: Presynaptic Ca^2+^ Spiking and Native Neuropeptide Release Peak As Somatic Ca^2+^ Declines

**DOI:** 10.1101/2023.12.01.569590

**Authors:** Markus K. Klose, Junghun Kim, Sydney N. Gregg, Brigitte F. Schmidt, Xiju Xia, Yulong Li, Edwin S. Levitan

**Affiliations:** Department of Pharmacology and Chemical Biology, University of Pittsburgh, Pittsburgh, PA 15261, USA; Department of Chemistry, Carnegie Mellon University, Pittsburgh, PA, 15213, USA; State Key Laboratory of Membrane Biology, School of Life Sciences, Peking University, Beijing 100871, China; PKU-IDG/McGovern Institute for Brain Research, Beijing 100871, China

**Keywords:** presynaptic Ca^2+^, GRAB sensor, peptidergic neurotransmission, circadian, sleep rhythms

## Abstract

*Drosophila* sLNv clock neurons release the co-packaged neuropeptides PDF and sNPF to regulate circadian behaviors (e.g., morning anticipation) and nighttime sleep^1-3^. Previous studies of membrane potential and cytoplasmic Ca^2+^ at the soma suggested that sLNv neuron activity peaks at night^4,5^, but exocytosis of neuropeptide-containing dense-core vesicles (DCVs) at their terminals peaks hours later at midmorning^6^. To resolve the basis of the timing mismatch between somatic physiology and terminal exocytosis, recently developed probes were used to measure daily rhythms in sLNv neuron synaptic Ca^2+^ and sNPF release. Remarkably, at midmorning after soma Ca^2+^ has dropped, both Ca^2+^ spiking and clock-dependent native neuropeptide release peak in the distal terminals of the protocerebrum. Furthermore, Ca^2+^ in the soma and terminals differ in dependence on Ca^2+^ influx. Finally, synaptic DCV exocytosis requires Ca^2+^ spike activity at terminals that is not evident at the soma. These results lead to two striking conclusions. First, soma Ca^2+^ recording, which is the focus of many circuit studies, is not indicative of presynaptic Ca^2+^ and neuropeptide release in distal sLNv terminals. Second, daily clock- and activity-dependent sLNv terminal neuropeptide release occurs ∼9-18 hours in advance of known sLNv neuropeptide effects on nighttime sleep and morning behavior.

## Results and Discussion

### Native synaptic neuropeptide release by sLNv neurons occurs at midmorning

Elevations in sLNv soma Ca^2+^ and resting membrane potential, which have been presumed to indicate when electrical activity and synaptic transmission occur, rise together at night^4,5^. However, synaptic exocytosis of DCVs assayed with an exogenous neuropeptide tagged with a fluorogen-activating protein (Dilp2-FAP)^7^, peaks hours later^6^. This timing discrepancy might arise because the latter method does not accurately quantify the exocytosis of native neuropeptides from DCVs, which in sLNv terminals contain PDF and sNPF. Therefore, we set out to measure endogenous neuropeptide release by sLNv terminals based on two recent developments. First, PDF and sNPF were found to be co-packaged together in the same individual DCVs in sLNv neurons^3^. This implies that assaying release of either neuropeptide would be sufficient to test if exocytosis timing matched native neuropeptide release. Second, while no sensor for PDF is available, a GPCR-activation-based sensor for detecting sNPF (GRAB_sNPF1.0_) was developed^8^. Therefore, GRAB_sNPF1.0_ expression was driven with PDF-GAL4 and sLNv neuron terminals, where neuropeptide-containing DCVs traffic to undergo exocytosis, were imaged for evidence of native neuropeptide release in brain explants.

Validation experiments included demonstrating dose-dependent responses to *Drosophila* sNPF2 (Fig. 1A). We then tested for a daily rhythm in GRAB_sNPF1.0_ signals at sLNv terminals. These experiments showed that the peak sNPF signal occurs at ZT 3 (Fig. 1B), which coincides with DCV exocytosis measured previously^6^. The sNPF peak cannot be attributed to a change in sensor expression because Fmax values do not change between ZT 23 and ZT 3 (Fig. 1C). Furthermore, selective knockdown of sNPF expression in sLNv neurons by RNA interference (RNAi) reduced GRAB_sNPF1.0_ fluorescence at ZT 3 (Fig. 1D). Thus, daily peaks in sNPF sensor fluorescence are the result of neuropeptide release from sLNv neurons rather than other sNPF-expressing neurons in the brain (Fig. 1D). Also, application of 100 µM Cd^2+^, which blocks Ca^2+^ channels, inhibited the release peak at ZT 3 (Fig. 1E). Therefore, Ca^2+^ influx is required for rhythmic synaptic neuropeptide release. Finally, consistent with the control of daily neuropeptide rhythms by the circadian clock^9^, the elevated sNPF sensor signal at ZT 3 was lost in the *per*^*01*^ clock gene amorphic mutant (Fig. 1F). Together, the above experiments show that Ca^2+^ influx-evoked endogenous neuropeptide release by sLNv terminals is triggered by the clock at midmorning.

**Figure 1.**
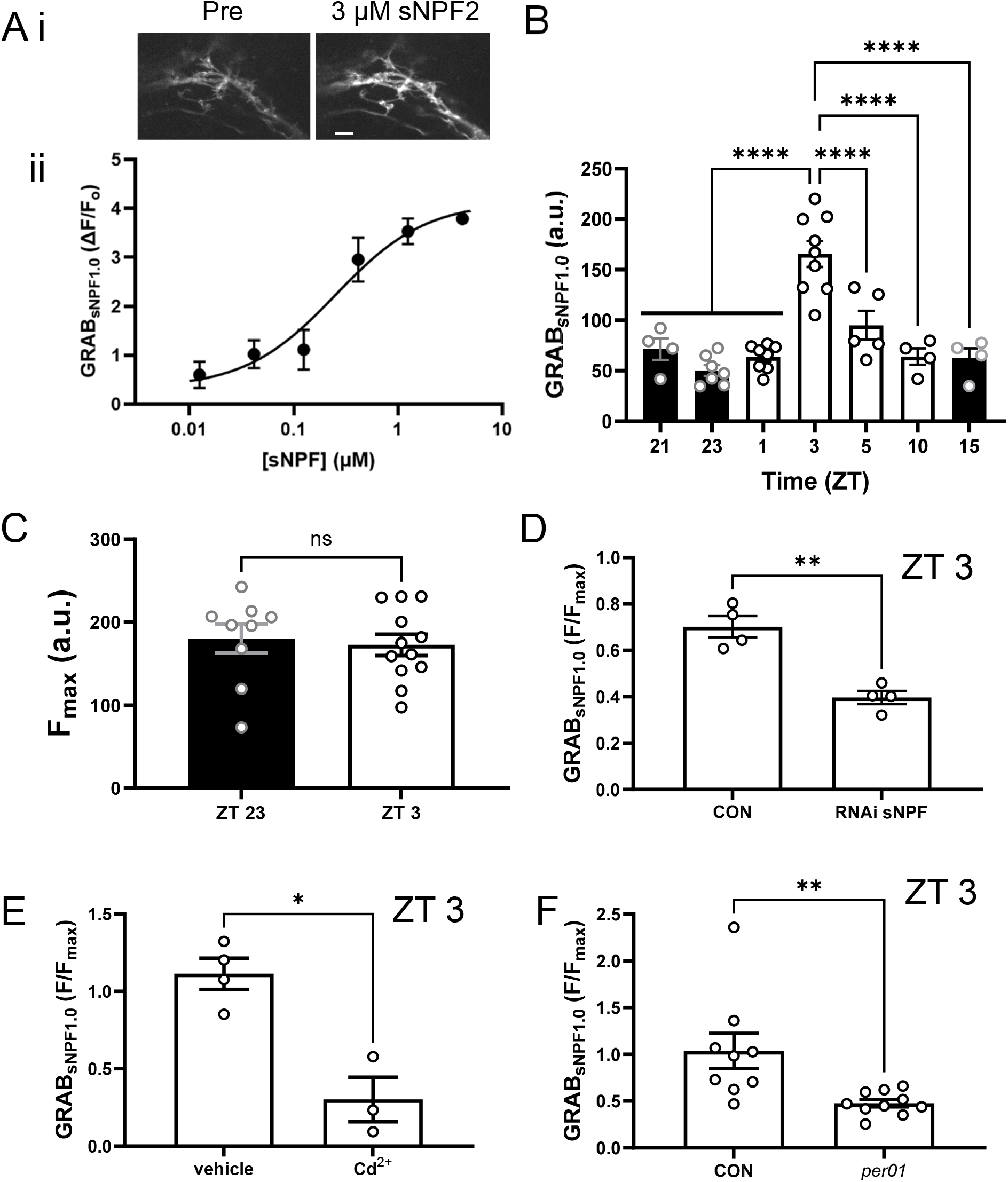
Circadian rhythm of native neuropeptide release from sLNv nerve terminals. A. (i) GRAB_sNPF1.0_ images in sLNv nerve terminals before (Pre) and after application of exogenous 3 µM sNPF2 to the brain explant. Shown images are maximum z-projections from image stacks made from 8 1 um steps. (ii) Dose-response curve for GRAB_sNPF1.0_ in LNv neurons in response to various concentrations of sNPF2. Note that responses are expressed in fold change relative to initial fluorescence (F_0_). Error bars here and in subsequent panels show standard error of the mean (SEM). B. Graph represents average GRAB_sNPF1.0_ fluorescence from sLNv nerve terminals at different times of day (ZT) in flies entrained to a 12 hour light:12 hour dark cycle. One-way ANOVA revealed a significant difference (P < 0.0001). Post-test analysis by Tukey’s multiple-comparison test is presented: ^****^ P < 0.0001, ^***^ P < 0.001. C. The maximal GRAB_sNPF1.0_ response evoked by 3 μM (F_max_) did not change between ZT 23 and ZT 3. D. GRAB_sNPF1.0_ fluorescence at peak release time of day (ZT3) in the sLNv terminals of control UAS-GRAB_sNPF1.0_; PDF-GAL4 flies (CON) and UAS-sNPF RNAi/UAS-GRAB_sNPF1.0_; PDF-GAL4 flies (RNAi sNPF) (t-test, P < 0.01). E. GRAB_sNPF1.0_ fluorescence at peak release time of day (ZT3) in the sLNv terminals 60 minutes after the application of vehicle or ∼100 µM cadmium chloride (Cd^2+^) (t-test, P < 0.05). F. GRAB_sNPF1.0_ fluorescence at peak release time of day (ZT3) in the sLNv terminals in controls and *per*^*01*^ (*per01*) flies (t-test, P < 0.01).

### Different Ca^2+^ rhythms in the sLNv soma and terminals

Because the shared timing of native neuropeptide release (Fig. 1) and DCV exocytosis^6^ does not correspond with reported elevations in soma Ca^2+^ and membrane potential^4,5^, we measured Ca^2+^ in terminals where synaptic neuropeptide release occurs with cytoplasmic GCaMP8f (GC) or synaptotagmin-fused mScarlet3-tagged GCaMP8f (ssGC), which enables ratiometric measurements. In the morning, sLNv terminal imaging of either GCaMP8f version revealed spontaneous presynaptic Ca^2+^ spikes that display a rapid rise and subsequent slower decay (Fig. 2Ai,ii and 2Bi), which is expected for presynaptic responses to action potentials^10,11^. Also consistent with activity responses, inhibiting Ca^2+^ influx with Cd^2+^ application blocked presynaptic Ca^2+^ spike activity within 5 minutes in every experiment (Fig. 2Aii-v, N = 12). We then determined the daily timing of spontaneous Ca^2+^ spike activity by imaging ssGC. Strikingly, the frequency of activity varies across the day with a peak of 1.8 ± 0.2 Hz at ZT 3 (Fig. 2B), which corresponds to the timing of peak synaptic neuropeptide release (Fig. 1B) and DCV exocytosis^6^.

**Figure 2.**
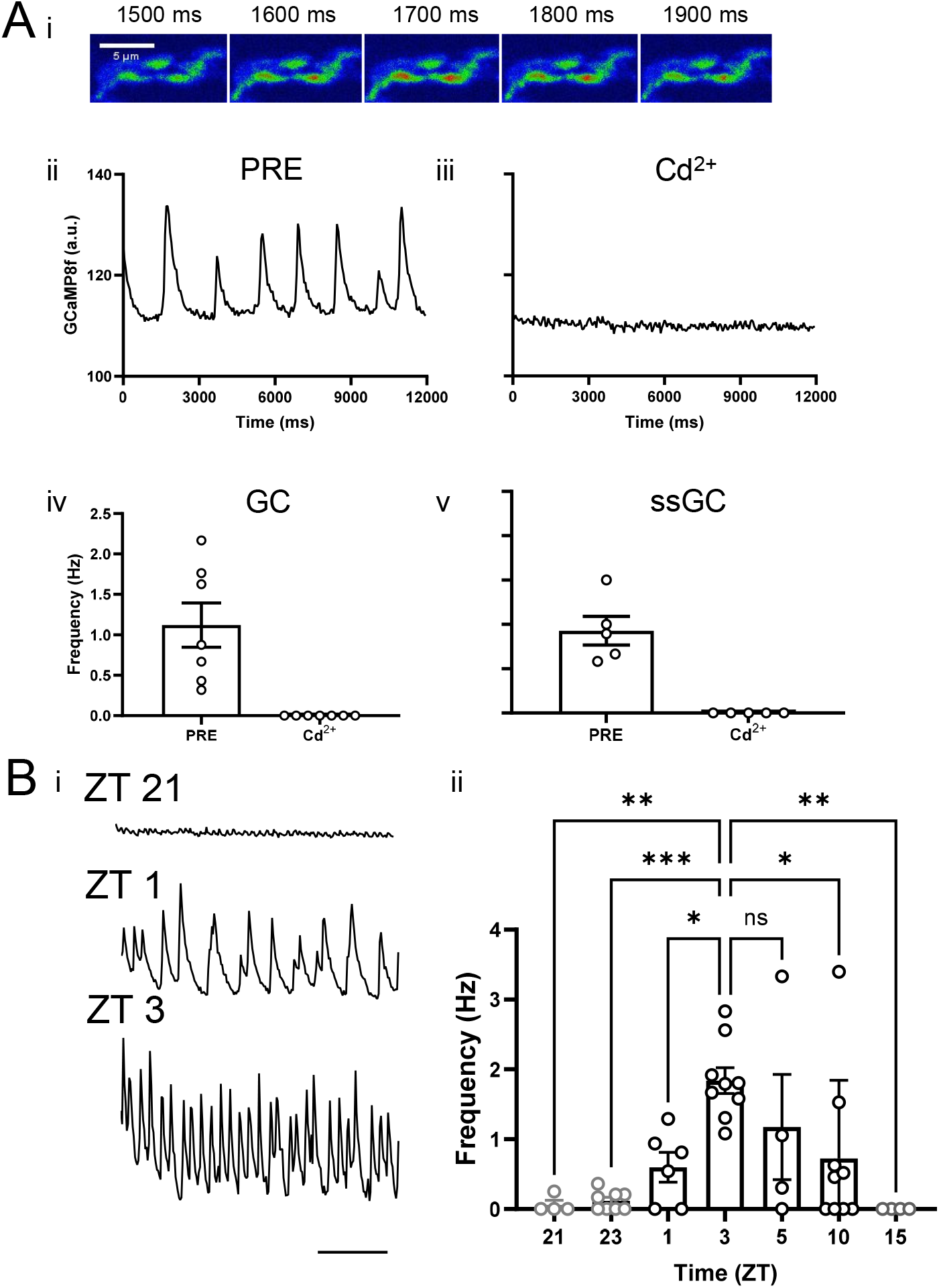
Presynaptic Ca^2+^ imaging in sLNv nerve terminals. A. (i) Cytoplasmic GCaMP8F driven by PDF-GAL4 in distal sLNv nerve terminals imaged over time at ZT 1. (ii) GCaMP8F fluorescence from one ROI measured at ∼20 Hz in normal HL3 saline. (iii) Graph represents GCaMP8F fluorescence from one ROI in HL3 saline before (PRE) and after adding ∼100 μM cadmium chloride (Cd^2+^). (iv) Morning (ZT1-5) activity frequency of sLNv nerve terminals expressing GCaMP8F before and after Cd^2+^ (∼100 μM). (v) Morning (ZT1-5) activity frequency of sLNv nerve terminals expressing Syt-mScarlet3-GCaMP8F (ssGC) before and after Cd^2+^ (∼100 μM). Note that ZTs were not matched between iv and v. B. (i) Syt-mScarlet3-GCaMP8F traces in distal sLNv nerve terminals imaged over time at ZT 21, 1 and 3. Bar indicates 3 s. (ii) Average Ca^2+^ transient frequencies measured in sLNv nerve terminals at different times of day using Syt-mScarlet3-GCaMP8F. One-way ANOVA revealed a significant difference (P < 0.0001). Post-test analysis by Tukey’s multiple-comparison test is presented: ^****^ P < 0.0001, ^***^ P < 0.001

The phase disparity between presynaptic Ca^2+^ reported here compared to soma Ca^2+^ reported previously^5^ could result from different experimental conditions or be the result of an unexpected disconnect between Ca^2+^ in the soma and terminals. To distinguish between these possibilities, soma Ca^2+^ was measured by ratiometric imaging of ssGC. Measurements in our preparations showed that soma Ca^2+^ drops during the morning (Fig. 3A, p < 0.0001 for least squares fit compared to a slope of 0) in agreement with a prior report^5^. Therefore, experimental conditions do not explain the soma-terminal Ca^2+^ disparity and peak Ca^2+^ firing frequency in terminals surprisingly occurs while soma Ca^2+^ is dropping.

**Figure 3.**
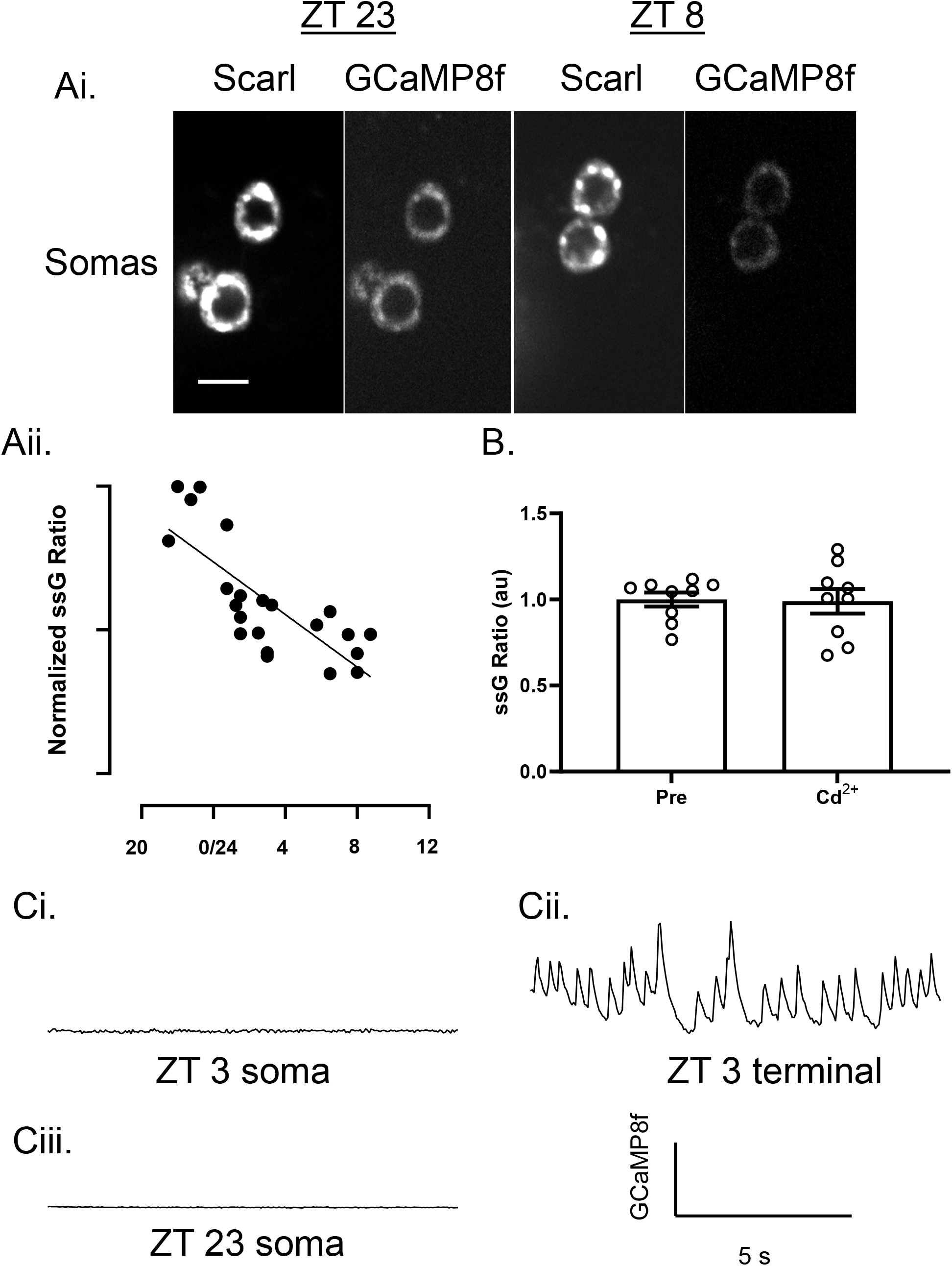
Somatic Ca^2+^ regulation in sLNv neurons. A. (i) ssGC fluorescence images showing the mScarlet and GCaMP8f signals in sLNv somas at ZT 23 and ZT 8, (scale bar, 5 µm). (ii) GCaMP8f/mScarlet ratios were calculated (ssG Ratio) and graphed from ZT 21-ZT 9 revealing soma Ca^2+^ is elevated at night and subsequently decreases throughout the morning. B. Addition of ∼100 µM CdCl_2_ for 20 minutes at ZT 23 had no effect on the soma ssG Ratio, revealing the elevated Ca^2+^ levels at this time were not the result of Ca^2+^ influx. C. At ZT 3, when terminals display maximal Ca^2+^ spike frequency, somatic recordings revealed no Ca^2+^ spikes (18 sLNv somas from 9 hemi-brains). (i) Cytoplasmic GCaMP8f in sLNv somas reveals no activity, (ii) while sLNv terminals in the same hemibrain revealed Ca^2+^ spike activity (repeated in 4 brains). (iii) At ZT 23, no firing was observed in sLNv somas (8 sLNv somas from 4 hemi-brains).

The disparate Ca^2+^ timing in the soma and terminals reflects different mechanisms operating in the two neuronal compartments. First, while terminal Ca^2+^ spikes and neuropeptide release depend on Ca^2+^ influx (Figs. 1 and 2), elevated soma Ca^2+^ at ZT 23 measured by ratiometric ssGC imaging persists when Ca^2+^ influx is blocked with Cd^2+^ for 20 minutes (Fig. 3B), indicating that the somatic Ca^2+^ increase does not require Ca^2+^ influx, while terminal Ca^2+^ spikes do. Second, high speed (∼20 Hz) imaging of GC showed that sLNv somas did not produce Ca^2+^ spikes at either ZT 23 (N = 8 somas from 4 hemibrains) when soma Ca^2+^ is elevated or ZT 3 (N = 18 somas from 9 hemibrains) when terminal Ca^2+^ spiking is greatest (Fig. 3C). This result was also evident in experiments in which both compartments were studied in a single hemibrain (N = 4). Therefore, different mechanisms, each peaking at different times, govern daily Ca^2+^ elevations in the sLNv neuron soma and terminals. Together, the above data show that synaptic neuropeptide release by sLNv neurons is triggered by spikes of Ca^2+^ influx at midmorning, which contrasts with the spike-free elevation of somatic Ca^2+^ that occurs late at night and does not depend on Ca^2+^ influx.

### Peak synaptic activity depletes a large releasable neuropeptide pool

As synaptic neuropeptide release is limited to the peak of activity, we considered why lower frequency activity prior to the peak (e.g., at ZT 1) did not elicit neuropeptide release (Figs. 1B and 2B). First, peak activity might be needed for efficient release because of the superlinear dependence of release on Ca^2+^. Alternatively, the lower activity prior to the peak may not evoke synaptic release because recently accumulated neuropeptide that occurs in preparation for release^6,9^ is in DCVs that are not yet competent for release (i.e., because they require priming). To test the latter hypothesis, the DCV exocytosis response to K^+^-induced depolarization was measured with Dilp2-FAP imaging^7^ at ZT 1, a time point when synaptic neuropeptide content has already increased, but daily exocytosis has not yet been initiated^6^. Strikingly, DCV exocytosis in response to depolarization, which was measured with the membrane impermeant fluorogen MG-Tcarb, is robust at ZT 1 (Fig. 4, Hi K^+^). Furthermore, the release evoked by depolarization at ZT 1 is comparable to peak endogenous release at ZT 3^6^, suggesting that endogenous rhythmic release depletes the releasable pool. To relate release to the total presynaptic pool, a membrane permeant version of the fluorogen (MGnBu^12^) was applied to label presynaptic Dilp2-FAP. This showed that the releasable pool is about half of the total neuropeptide pool in terminals (Fig. 4, MGnBu). Thus, at ZT 1 there is a large pool of release-competent DCVs in sLNv terminals. Therefore, the clock-dependent timing of release is not due to a change in vesicle release competence, but instead reflects the frequency requirement for robust synaptic neuropeptide release and the timing of activity-dependent Ca^2+^ influx that occurs in terminals, but not the sLNv soma.

**Figure 4.**
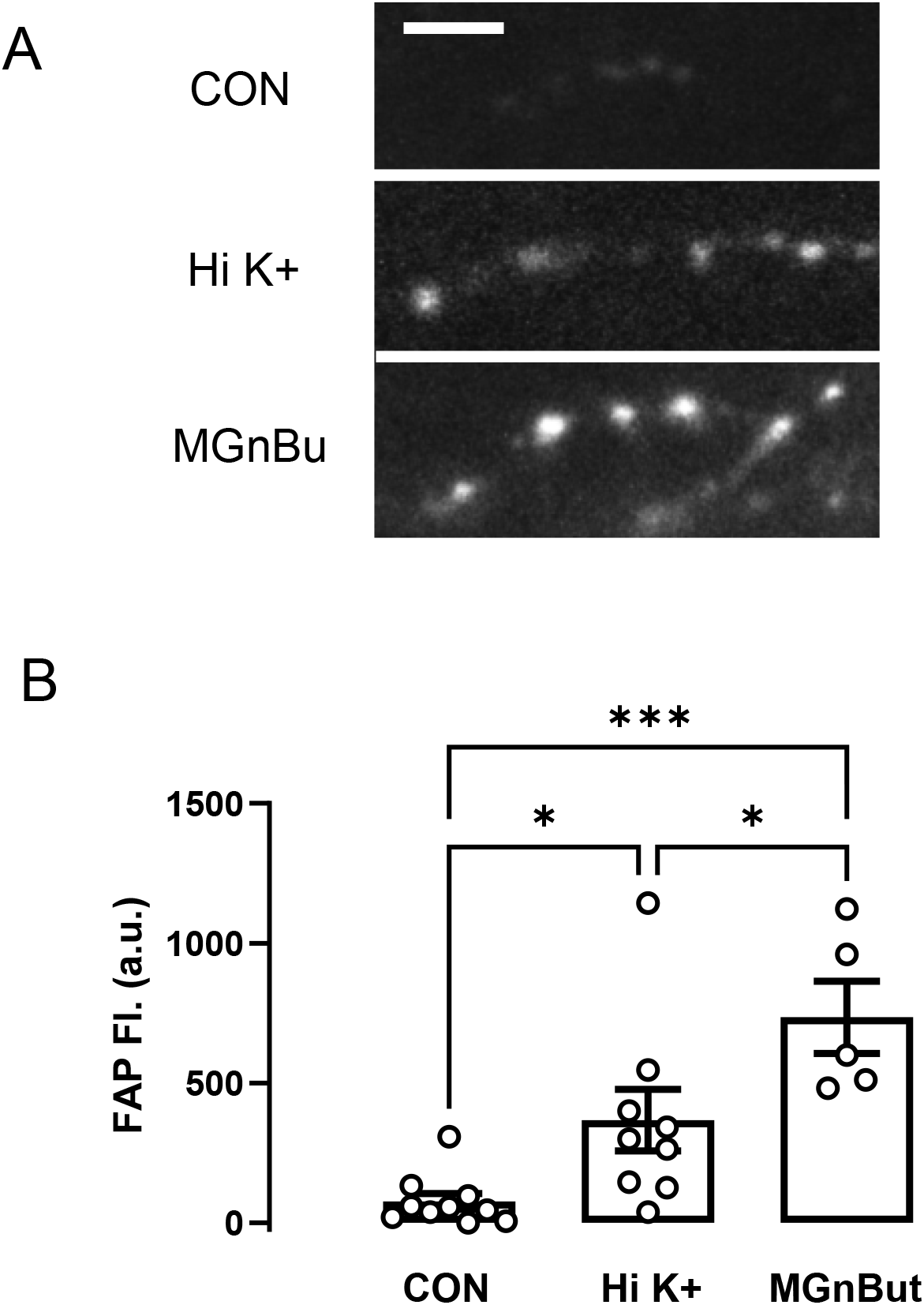
A large releasable pool of DCVs is present at ZT 1 in sLNv nerve terminals. A. Confocal images of the FAP signal in sLNv terminals following one-hour exposure to MG-Tcarb in normal HL3 saline (CON), MG-Tcarb in high K^+^ saline (Hi K+), or with exposure to the membrane permeant MG analog MGnBu (500 nM) in normal HL3 saline. Scale bar is 10 µm. B. Quantification of FAP-MG fluorescence in sLNv nerve terminals exposed to MG-Tcarb in normal HL3 saline (CON), MG-Tcarb in Hi K+ HL3 saline, or membrane permeable MGnBu in normal HL3 saline. ^***^ P < 0.001, ^*^ P < 0.05. n = 10 for CON; n = 9 for Hi K+; n = 6 for MGnBu. One-way ANOVA revealed a significant difference (P < 0.001). Post-test analysis by Tukey’s multiple-comparison test is presented: ^***^ P < 0.001, ^*^ P < 0.05.

## Conclusions

The first conclusion derived from this study is that soma Ca^2+^ recordings, which are often used to infer the timing of neuronal activity, do not always reflect the timing of activity and neurotransmission. For sLNv clock neurons, rhythmic Ca^2+^ elevations in the soma occur without Ca^2+^ influx many hours before presynaptic Ca^2+^ influx-dependent activity and neuropeptide release at terminals. Therefore, the timing of peptidergic transmission by sLNv terminals is not evident from soma Ca^2+^. These results show circuit activity cannot be deduced from soma Ca^2+^ imaging with certainty. Thus, circuit models that were formulated based solely on soma Ca^2+^ imaging should be re-evaluated taking into account the potential for uncoupling between the soma and terminals, which was demonstrated here with a clock neuron.

The second major finding of this study is that rhythmic activity-dependent native neuropeptide release from sLNv synapses occurs at midmorning. Therefore, synaptic sNPF release occurs >9 hours before nighttime sleep, which is promoted by sNPF from sLNv neurons^2^. Also, because synaptic PDF is co-packaged with sNPF^3^ and its release from sLNv neurons is important for morning anticipation just prior to sunrise^1^, the rhythmic synaptic release demonstrated here occurs >18 hours in advance of a PDF-dependent behavior. The extraordinary time delays between neuropeptide release by sLNv neurons and neuropeptide-elicited behavioral effects raises fundamental questions about how peptidergic circuits control rhythmic circadian and sleep behaviors.

## Materials and methods

### Physiology

All flies were entrained for at least for 72 hours in a 12-hour light: 12-hour dark (LD) schedule before dissection of 4 to 9 day-old males to generate brain explants. Dissections during the dark phase were performed under a red light. Adult flies were immobilized with CO_2_ gas and brains were dissected in 0 Ca^2+^ HL3 saline solution (70 mM NaCl, 5 mM KCl, 20 mM MgCl_2_*6H_2_O, 115 mM Sucrose, 5 mM Trehalose, 5 mM Hepes, and 10 mM NaHCO_3_, pH 7.3) and then put into polylysine-coated plastic dishes containing HL3 with 2 mM Ca^2+^ for imaging^13^. High potassium (Hi K+) saline was used to elicit neuropeptide release (35 mM NaCl, 80 mM KCl, 10 mM MgCl_2_^*6H^_2_O, 65 mM Sucrose, 5 mM Trehalose, 5 mM Hepes, and 10 mM NaHCO_3_, pH 7.3).

All imaging was done on setups with upright Olympus microscopes equipped with a 60× 1.1 NA dipping water immersion objective, a Yokogawa spinning disk confocal head, lasers (i.e., 488 nm laser for GFP illumination, a 561 nm laser for mScarlet3 detection, a 640 nm laser for FAP imaging), and a Teledyne Photometrics sCMOS camera.

Two fluorescent probes were expressed using the PDF-GAL4 driver to image neuropeptide release from sLNv terminals is the dorsal protocerebrum. The first was the sNPF sensor, GRAB_sNPF1.0_^8^, and the other was Dilp2-FAP^7^, which detects DCV fusion pore opening events. UAS-GRAB_sNPF1.0_ expression in sLNv terminals was used to detect both sNPF endogenous release and application of exogenous sNPF2 (WFGDVNQKPIRSPSLRLRF) (GenScript Life Science). FAP imaging experiments were performed as previously described^6^. In the current experiments, a recombinant with UAS-Dilp2-GFP^14^ and UAS-Dilp2-FAP was used so that GFP could be used for focusing before application of the fluorogens MG-Tcarb (membrane impermeant) or MGnBu (membrane permeant)^12^. Ca^2+^ was imaged with UAS-GCaMP8f (GC) or UAS-syt-mScarlet3-GCaMP8f (ssGC) driven by PDF-GAL4. For resolving Ca^2+^ spikes, the green channel data were streamed with 50 ms exposures. ssGC ratios were based on alternating between GFP and red optics (e.g., 488 nm excitation for GCaMP and 561 nm excitation of mScarlet) with 100 ms exposures for each channel recorded at 2 Hz, while FAP imaging used Cy5 far red optics (640 nm excitation). Quantification of fluorescence was done in Imagej or Fiji. Statistical analysis (e.g., tests and calculation of standard error of the mean (SEM) for error bars) was performed with Graphpad Prism software.

### Fly lines

All flies used the PDF-Gal4 promoter on the third chromosome (provided by Paul Taghert, Washington University in St. Louis). PDF-GAL4 drives expression in the two subsets of clock neuron that express PDF neuropeptide, the small ventrolateral (sLNv) neurons and the large ventrolateral (lLNv) neurons, of which only the sLNv neurons express sNPF. UAS-Dilp2-GFP, UAS-Dilp2-FAP, UAS-GRAB_sNPF1.0_ and UAS-synaptotagmin-mScarlet3-GCaMP8f flies, with the latter coming from Dion Dickman (University of Southern California), were reported previously^7,8,11,14^. *w*^*1118*^ flies were from Zachary Freyberg (University of Pittsburgh), while UAS-sNPF RNAi (III) flies were from Leslie Griffith (Brandies University). Lines from the Bloomington *Drosophila* stock center included *Bl#* 92857 (UAS-GCaMP8f) and *Bl#* 80917 (*per*^*01*^).

## >Acknowledgments

We thank Dion Dickman and Leslie Griffith for flies. Stocks obtained from the Bloomington Drosophila Stock Center (NIH P40OD018537) were used in this study. We are also thankful for suggestions from Marcel Bruchez (Carnegie Mellon University, deceased) and comments on the manuscript from David Deitcher (Cornell University).

## References

1. Crespo-Flores SL, Barber AF. The Drosophila circadian clock circuit is a nonhierarchical network of peptidergic oscillators. Curr Opin Insect Sci. 2022; 52:100944.

2. Shang Y, Donelson NC, Vecsey CG, Guo F, Rosbash M, Griffith LC. Short neuropeptide F is a sleep-promoting inhibitory modulator. Neuron. 2013;80(1):171–83.

3. Yu J, Zhang Y, Clements K, Chen N, Griffith LC. Genetically-encoded markers for confocal visualization of single dense core vesicles. Commun Biol. 2025;8(1):383.

4. Cao G, Nitabach MN. Circadian control of membrane excitability in Drosophila melanogaster lateral ventral clock neurons. J Neurosci. 2008;28(25):6493–501.

5. Liang X, Holy TE, Taghert PH. Synchronous Drosophila circadian pacemakers display nonsynchronous Ca^2+^ rhythms in vivo. Science. 2016; 351(6276):976–81.

6. Klose MK, Bruchez MP, Deitcher DL, Levitan ES. Temporally and spatially partitioned neuropeptide release from individual clock neurons. Proc Natl Acad Sci U S A. 2021;118(17):e2101818118.

7. Bulgari et al. Activity-evoked and spontaneous opening of synaptic fusion pores. Proc Natl Acad Sci U S A. 2019;116(34):17039–17044.

8. Xia X, Li, Y. A new GRAB sensor reveals differences in the dynamics and molecular regulation between neuropeptide and neurotransmitter release bioRxiv. 2024; doi: 10.1101/2024.05.22.595424.

9. Park JH, Helfrich-Förster C, Lee G, Liu L, Rosbash M, Hall JC. Differential regulation of circadian pacemaker output by separate clock genes in Drosophila. Proc Natl Acad Sci U S A. 2000;97(7):3608–13.

10. Macleod GT, Hegström-Wojtowicz M, Charlton MP, Atwood HL. Fast calcium signals in Drosophila motor neuron terminals. J Neurophysiol. 2002;88(5):2659–63.

11. Li X, Chien C, Han Y, Sun Z, Chen X, Dickman D. Autocrine inhibition by a glutamate-gated chloride channel mediates presynaptic homeostatic depression. Sci Adv. 2021;7(49):eabj1215.

12. Perkins LA, Fisher GW, Naganbabu M, Schmidt BF, Mun, F, Bruchez MP. High-content surface and total expression siRNA kinase library screen with vx-809 treatment reveals kinase targets that enhance F508del-CFTR rescue. Mol Pharm 2018; 5(3):759–767.

13. Klose M, Duvall L, Li W, Liang X, Ren C, Steinbach JH, Taghert PH. Functional PDF Signaling in the Drosophila Circadian Neural Circuit Is Gated by Ral A-Dependent Modulation. Neuron. 2016;90(4):781–794.

14. Wong MY, Zhou C, Shakiryanova D, Lloyd TE, Deitcher DL, Levitan ES. Neuropeptide delivery to synapses by long-range vesicle circulation and sporadic capture. Cell. 2012;148(5):1029–38.

